# Characterization of a Novel *Tequatrovirus* Phage from Pristine Stretch of The Ganges River, India, in Reducing Bacterial Load from Sewage Water

**DOI:** 10.1101/2024.11.01.621489

**Authors:** Rachel Samson, Ameya Pawar, Krishna Khairnar, Mahesh Dharne

## Abstract

Effective treatment of wastewater (WW) and its reuse is necessary to meet certain sustainable development goals and a circular economy. *Escherichia coli* is one of the primary contaminants in the WW, and its extra-intestinal occurrence poses a considerable threat under one health. This study is the first report of a novel broad-spectrum phage (фERS-1) isolated from a pristine stretch of the Ganges River in the biocontrol of *E. coli*, resistant to 3^rd^-and 4^th^ generation cephalosporins and aztreonam. This is the first report of a phage from the *Tequatrovirus* genus to infect *P. aeruginosa*. The фERS-1 could reduce the abundance of *E. coli* cells by 8.22 log_10_ CFU/mL ≤24 hrs. Additionally, □ ERS-1 disrupted the biofilm of *E. coli* with a reduction of 3.88 log_10_ CFU/mL. Further, □ ERS-1 could inhibit biofilm by multiple strains of *E. coli* and multiple genera (*E. coli, S. boydii,* and *P. aeruginosa*). The phage □ ERS-1 reduced bacterial counts in raw WW by 2 log_10_ CFU/mL and 4 log_10_ CFU/mL reduction in coliform-enriched WW in ≤24 hours. Overall, this study suggests that □ ERS-1 could be used as an effective alternative to be combined with other treatments for improving the quality of WW disposal and environmental health by reducing the bacterial load.

**Graphical Abstract:** 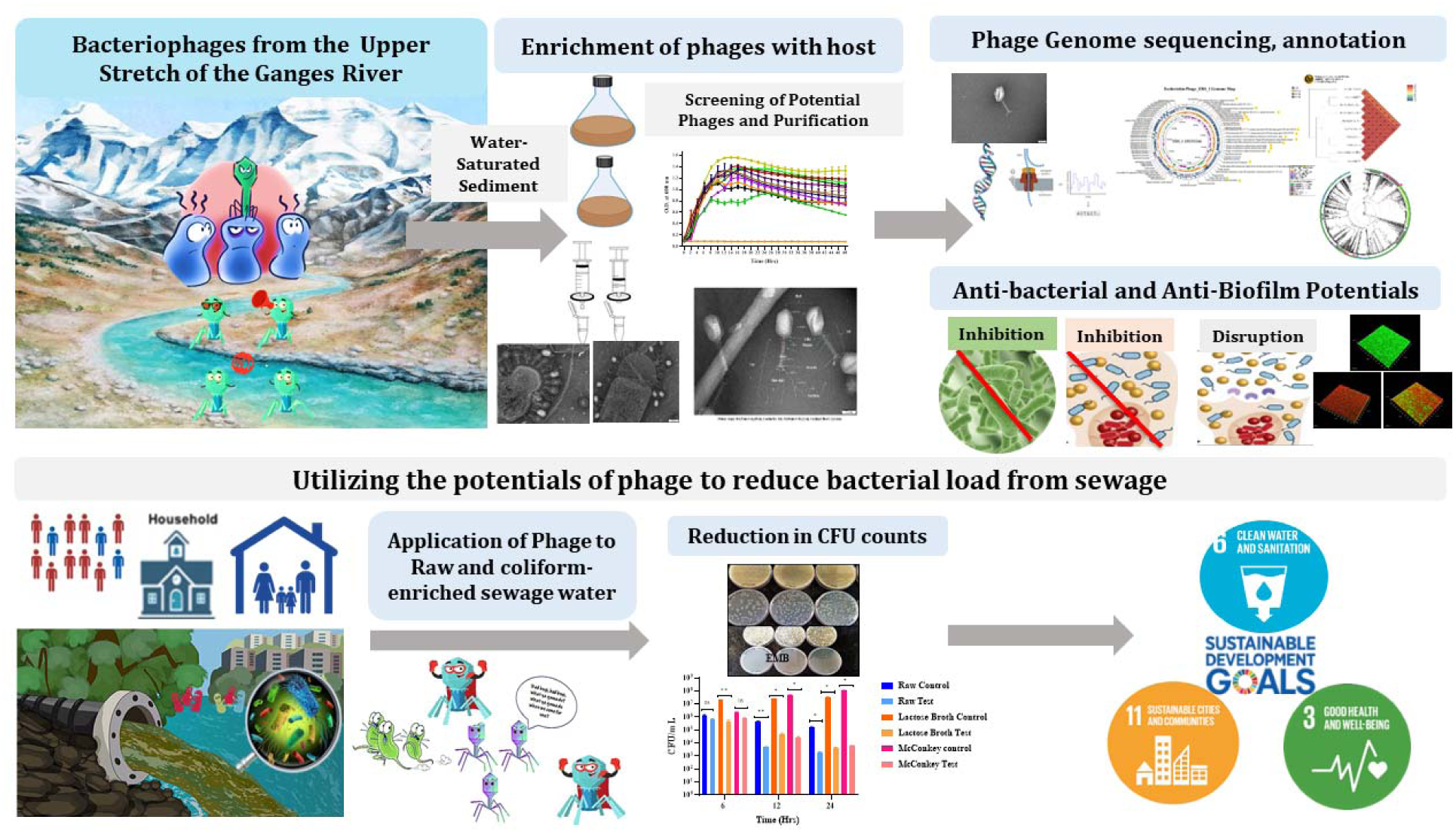

**Highlights:**

- Isolation of a novel phage from a pristine stretch of the Ganges River
- Antibiofilm activity against *E. coli* >8 log_10_ inhibition, >3 log_10_ disruption
- Biofilm inhibition of >50% against *P. aeruginosa* and *S. boydii*
- 2 log_10_ and 4 log_10_ reduction of bacterial counts in phage-treated raw sewage

## 1. Introduction

The treatment and recycling of wastewater (WW) for non-potable purposes has been recognized progressively as a sustainable alternative as it is a cost-effective option to tackle the existing water scarcity crisis. Reusing domestic WW allows sustainable use of water resources for agriculture or environmental benefits, contributing to the circular economy (Hernández-Crespo et al., 2022). Considering the load of inorganic and organic discharges in the sewage, the conventional indicators include estimating the carbon pollutants and nitrogen and phosphorus content. However, in recent times, biological indicators have been at the centre of attention for sewage treatment and reuse (Al-Gheethi et al., 2018).

The raw (untreated) WW comprises various pathogenic bacteria, which include the members of *Enterobacter* sp., *Staphylococcus aureus, Klebsiella* sp*., Acinetobacter* sp., *Escherichia coli, Enterococcus* sp., *Proteus* sp., *Salmonella* sp., *Shigella* sp., *Pseudomonas aeruginosa,* and *Citrobacter* sp. Such microbial contamination in water has a detrimental effect on the public health (Soliman et al., 2023). The key bacterial pathogens in sewage that cause environmental and health problems are *E. coli, Salmonella, Shigella,* and *Pseudomonas*. The contamination of these microbes in the environment occurs through intestinal and extra-intestinal routes (Jang et al., 2017). Therefore, biological indicators should be more focused on reflecting the connotation of sewage recycling better and providing an answer to the current water-environmental sanitation practices. The current WW treatment includes a combination of physical, chemical, and biological methods for eliminating microbial contamination. (Qian et al., 2022). However, the rise of antibiotic resistance in bacteria has considerably challenged the treatment of microbial pollution in WW. Therefore, to tackle the existing antibiotic resistance crisis, there is a need to safeguard human and animal health by shielding environmental health through the ‘one health’ approach (Garvey, 2020).

*E. coli* members are an essential indicator in evaluating the extent of environmental pollution. Additionally, *E. coli* has been identified as a key enteric pathogen responsible for diarrhoeal disease due to its transmission through contaminated food, water, soil, surfaces, and hands (Navab-Daneshmand et al., 2018). The fate of *E. coli* strains as commensals or pathogens (expressing virulence factors) depends upon a complex balance between the host’s status and expression of the virulence determinants. Certain signature characteristics of pathogenic strains of *E. coli* include adhesion, biofilm formation, toxin production, and evasion of host defense mechanisms. The environmental transmission of *E. coli* and its pathotypes include animal wastes, manure, WW, and sewage sludge exiled from WW treatment plants (Osińska et al., 2023). Besides, the presence of *E. coli* in treated WW remains a considerable public health concern due to the high expense of removal via traditional techniques employed in sewage treatment plants. Therefore, to address the issue of AMR and control microbial contamination, bacteriophages as green biocontrol agents have re-emerged (Soliman et al., 2023).

Recent studies have highlighted the extended ability of phages to control bacterial populations in various fields of agriculture, aquaculture, biomedicine, the food industry, and wastewater treatment. However, there are limited studies detailing the use of cocktail of phages in their natural or engineered form to treat the WW for the reduction of bacterial load and the elimination of waterborne pathogenic bacteria (Beheshti Maal et al., 2015; Bhargava et al., 2023; Grami et al., 2022; Jassim et al., 2016; Periasamy and Sundaram, 2013; Withey et al., 2005). Most of the phages are host-specific and facilitate the killing of their host through lysis, followed by the release of the phage progeny. Moreover, phages, being abundant in nature, make them ideal candidates for their use in a variety of applications spanning various domains under one health canopy (Samson et al., 2024). However, a major limitation of the use of phages in environmental settings is their narrow host range. Isolation of phages that could lyse multiple hosts effectively is challenging. However, this constraint is overcome by combining various monovalent phages as a cocktail or engineering naturally occurring phages to extend their host spectra. Combining different phages may often lead to antagonistic outcomes and cause a bacteriostatic effect instead of bactericidal ones (Zhou et al., 2022).

The Ganges River is India’s national river and is well known for its unique properties of ‘self-cleansing (non-putrefying) and special healing’ since the ancient past (Khairnar, 2016). The waters of the River Ganges have a rich history of demonstrating antibacterial properties, first studied by Ernst Hankin against *Vibrio cholerae* in 1896 (Hankin, 1896). However, a detailed insight into these antibacterial aspects was provided by Félix d’Hérelle, who discovered and termed the antagonist microbe of *V. cholerae* as ‘bacteriophage’ (D’Herelle, 1917).

It is well-known that despite tremendous anthropogenic activities and their related pollution load, the Ganges can rejuvenate itself rapidly (Paul, 2017; Reddy and Dubey, 2019; Samson et al., 2019; Zhang et al., 2018). This fact has also been witnessed recently during COVID-19 lockdown times with remarkably clean water even at the most polluted sites (Dutta et al., 2020). However, there are limited studies with this riverine system wherein novel virulent phages were isolated from its waters against MDR strains of *K. pneumoniae* (Sundaramoorthy et al., 2021) and *P. aeruginosa* (Rathor et al., 2022). Our recent study on the sediments of the river Ganges along its 1500 km has identified the repertoire of bacteriophages and their associated host-phage functions against putative human, plant, and putrefying pathogens (Samson et al., 2023). Therefore, this unique aquatic ecosystem provides an opportunity to bioprospect its untapped and unique phage diversity for its potential applications. Given this, the present study was initiated to explore the untapped phage diversity from the pristine stretch of the river Ganges to isolate novel phages and explore their potential for use as biocontrol agents against *E. coli*, facilitating improved environmental health.

## 2. Materials and Methods

### 2.1. Bacterial strain and antimicrobial susceptibility profile

*Escherichia coli* (ATCC 8739) was the bacterial host used in this study. The genomic identity of the isolate was confirmed using the MinION Mk1C Nanopore sequencer (Oxford Nanopore Technologies). The resistance profile of the isolate was ascertained with the VITEK 2 system (bioMérieux) using two different sets of antimicrobial susceptibility testing (AST) cards, AST-N235 and AST-N281, having a defined set of parameters as recommended by the Clinical and Laboratory Standards Institute (CLSI) **(Supporting Information, *E. coli_*AST_281)**.

### 2.2. Isolation, enrichment, and propagation of *Escherichia coli* phage □ ERS-1

Water-saturated sediment samples of the Ganges River from the Harshil (31°02’18.0“N 78°44’22.7” E) location were used as the source of phage isolation against the primary host *E. coli* (ATCC 8739). A different approach was used for the isolation of bacteriophages. The sediment samples (n=3, 50 gm) were vortexed at maximum speed for 30 minutes at room temperature. The sediment was allowed to settle briefly while the sediment-laden water (∼40 mL) was equilibrated with 10 mL of SM buffer for 1 hour in an incubator at 37□ with a speed of 180 rpm. The resulting mixture was removed and centrifuged (Eppendorf centrifuge, 5804 R) at 6000×g for 10 mins. The supernatant (∼40 mL) was then passed through a combination of 0.45μm 0.22μm of Polyether sulfone (PES) membrane-based syringe filters (Hi-Media), respectively, and 10 mL of filtrate from each set of the syringe filters was used for enrichment.

A total of 5mL of double-strength broth of soybean-casein digest broth (SCDB) (MH011-500G, Hi-Media) and 5 mL of exponential phase culture of *E. coli* (host) was added to 20 mL of the filtrate obtained from the previous step. The enrichment flasks were incubated at 37°C for 24 h with shaking (180rpm). The presence of lytic phages in the samples was confirmed by observing clear zones (plaques) against the bacterial lawn through the qualitative spot assay. The quantitative enumeration of phages in the enriched lysate was done using the soft agar overlay method. A single, well-isolated plaque was picked and suspended in SM (Sodium-Magnesium) buffer (NaCl 5.8g/L, MgSO_4_.7H_2_O 2g/L, Tris-HCl (pH7.4) 50mL, autoclaved distilled water 950mL). The phage titer of the released phages from the single plaque was determined in triplicates for over four times by making serial dilutions and infecting the log-phase culture of *E. coli*, followed by double-layer agar plating and incubation at 37 °C for 16-18 h. Purification and concentration of □ ERS-1 was carried out with Amicon® Ultra-15 centrifugal filter with Ultracel100 kDa membrane (Merck Millipore) (Sun et al., 2014). The multiplicity of infection (MOI) (Kasman et al., 2002) and Poisson distribution for predicting the probable number of infected cells in a population (P_0_) (Abedon, 2023) were calculated using the following formula:

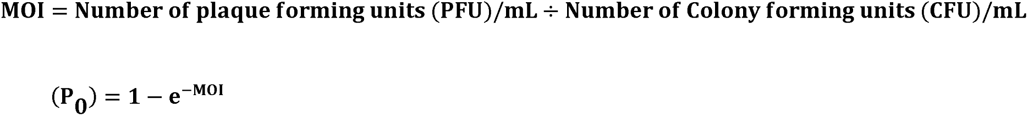

A high titer phage stock of the purified phage lysate containing 10^17^ PFU/mL was made using replicates of a total of 30 confluent lysis plates using replicates of a total of 30 confluent lysis plates flooded with 7mL of SM buffer and incubated at 15 with shaking (100 rpm) for 4 hours followed by ultracentrifugation at 14000×g for 30 mins at 4 and subsequent filtration of the lysate with 0.2 μm PES syringe filters (Bonilla and Barr, 2018). Enumeration of the high titer phage stock was done using the double-layer agar overlay method. The purified high-titer phage stock was stored at 4°C.

### 2.3. Characterization of *Escherichia coli* phage (□) ERS-1

#### 2.3.1 Morphological features

The morphology of □ ERS-1 was visualized using Jeol JEM-F2100 high resolution-transmission electron microscope (HR-TEM) at 200 kV and imaged with a Xarosa emsis camera coupled to the microscope. To obtain a detailed (3D) view of the phage morphology, HR-TEM was used in STEM (Scanning Transmission Electron Microscopy) mode, enabling scanning images at nanometer (nm) resolution. Briefly, 1 mL of the high titer stock of □ ERS-1 was centrifuged at 12,000 ×g for 30 mins, and the pellet was resuspended with 10 µL of SM buffer. A total of 4 µL of the sample was placed on a lacey formvar/carbon, 200 mesh, copper grid followed by staining with VitroEase™ Methylamine Tungstate Negative Stain (Thermo Fisher Scientific), following the manufacturer’s protocol. Morphological classification of phage was done following the guidelines recommended by the International Committee on Taxonomy of Viruses (ICTV) (Turner et al., 2023).

#### 2.3.2 Phage adsorption rate and one-step-growth curve

To understand how fast phage virions can adsorb a target bacterial host cell, an adsorption experiment was performed as described by (Heineman and Bull, 2007) with certain variations. The host cell culture at the exponential phase was mixed with the fresh phage lysate at an MOI of 0.1 and incubated at 37°C, 100 for 30 mins. After 5 minutes, an aliquot was drawn and centrifuged to pellet the adsorbed fraction of the phage each time. At the same time, the supernatant was filtered using a 0.2 μm PES syringe filter to obtain the fraction of unadsorbed phages. The fractions were plated using the agar overlay method to get total phage (N _total_) and free phage (N _free_) densities, respectively. The assay was carried out in three replicates, and the adsorption curve as a function of the percentage of adsorbed and unadsorbed phages versus time was plotted using GraphPad Prism v9.5.1. The adsorption rate (α) for each time point was calculated as:

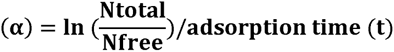

One one-step growth curve assay for □ ERS-1 was carried out as described by (Jagdale et al., 2019) with few modifications. Briefly, fresh phage lysate of □ ERS-1 at a MOI of 0.1 was added to the *E. coli* cells (10^8^ cells/mL) and allowed to adsorb for 20 mins at 37°C, 100 rpm. The adsorbed phages were centrifuged at 10,000 ×g for 5 min, and subsequently, the bacteria-phage complex (pellet) was resuspended in 2 mL of SCDB and incubated at 37°C, 100 rpm for 100 mins. A 100 µL aliquot was collected every 10 minutes, followed by plaque assay to determine the phage count (PFU/mL) at each time interval. The assay was carried out in triplicates.

#### 2.3.3 Evaluation of pH and temperature stability

The stability of □ ERS-1 under different acidic and basic conditions was assessed as described by (Oliveira et al., 2020). Briefly, the pH of the SM buffer was adjusted to 3, 5, 7, 9, and 11, followed by adding 100μL of the phage suspension (10^17^ PFU/mL) to 900 μL of SM buffer with respective pH. The samples were incubated for four hours at 37°C, 100 rpm in a dry bath, followed by plaque assay. For determining the thermal stability, suspensions of 100μL of the phage (10^17^ PFU/mL) in 900 μL of SM buffer were made and incubated at a diverse set of temperatures (4, 15, 25, 37, 45, 55, and 65°C) for four hours at 100 rpm followed by measuring the phage titer with plaque assay. The assays for pH and thermal stability of □ ERS-1 were carried out at two independent times in triplicates. GraphPad Prism v9.5.1 was used to compute the statistical significance of the results using repeated measures (RM) one-way analysis of variance (ANOVA). Additionally, multiple comparisons with a false discovery rate (FDR correction) and a p-value of 0.05 were used to compare the mean between the two groups.

#### 2.3.4 Evaluation of the Lytic Spectrum

The host range of □ ERS-1 was evaluated using in-house bacterial host cultures of *Escherichia coli* O157:H7 (ATCC 43888)*, Shigella boydii* (ATCC 9207, 8700)*, Pseudomonas aeruginosa* (ATCC 9027)*, Salmonella enterica* (ATCC 12011, 13314)*, Staphylococcus aureus* (ATCC 6538)*, Listeria monocytogenes* (ATCC 19112)*, Enterococcus faecalis* (ATCC 19443) *Yersinia enterocolitica* (ATCC 27729). Details of the susceptibility profile of each host strain with phage □ ERS-1 have been tabulated **(Table S1)**.

#### 2.3.5 DNA extraction, Genome Sequencing, and Analysis of **D** ERS-1

A high-titer phage stock was used to isolate DNA from phage □ ERS-1. The DNA was extracted using PureLink Viral RNA/DNA Mini Kit (Thermo Fisher Scientific), following the manufacturer’s protocol. A 16S rRNA gene amplification was done to ascertain that the extracted DNA was devoid of host DNA. The PCR products were examined on 0.8% agarose gel electrophoresis along with controls and standard molecular weight markers. The gel was visualized using the BioEra Gel Documentation System **(Fig S1)**.

The phage DNA concentration was quantified on a Qubit 4 Fluorometer (Invitrogen) using a dsDNA HS (High Sensitivity) assay kit fluorometer (Invitrogen). Library preparation of the samples for whole genome sequencing was carried out with Ligation Sequencing Kit (SQK-LSK114) and Native Barcoding Kit (SQK-NBD 114.24) as per the manufacturer’s instructions with certain modifications **(Text S1)**. The library was loaded onto flow cell R10.4.1(Oxford Nanopore Technologies), and the sequencing run was carried out for ∼56 hours using the MinION Mk1C sequencing platform.

The reads were processed from the sequencing data for quality control and adapter trimming using FastQC (Galaxy Version 0.25.1+Galaxy0) and Porechop (Galaxy Version 0.2.4+galaxy0). The genome assembly was performed using Flye, a *de novo* genome assembler v 2.9.1. Validation of sequence homology for the generated assembly with known phage sequences was done with NCBI nucleotide BLAST(BLASTN). Further, the phage genome annotation of the putative proteins was predicted with Rapid Annotation using the Subsystem Technology (RAST) annotation server (https://rast.nmpdr.org). The GenBank file generated from RAST was used in Proksee for an in-depth analysis and visualization of the phage genome (Grant et al., 2023).

Additionally, an open reading frame (ORF) finder from NCBI (https://www.ncbi.nlm.nih.gov/orffinder/) was used to identify the protein-coding sequences in the phage genome. Further, VipTree was used to understand genome-wide similarities between □ ERS-1 and other reference viral genomes (Nishimura et al., 2017). Whole genome Average nucleotide identity (ANI) to ascertain genetic relatedness between □ ERS-1 and other closely related phages was computed with OAT v0.93.1 (Orthologous Average Identity Tool) software (Lee et al., 2016). PhaBOX, a comprehensive web tool for phage identification, taxonomic classification, and prediction of phage lifestyle and its host, was used for an integrated phage analysis. Furthermore, intergenomic comparisons were made to ascertain relatedness between the phages using Virus Intergenomic Distance Calculator (VIRIDIC) (Moraru et al., 2020).

### 2.4. Antibiofilm Potentials of **D**ERS-1

#### 2.4.1 Quantification and Visualization of Biofilm Inhibition Spectrum of D □ ERS-1

A mixed culture biofilm inhibition assay was performed to understand □ ERS-1’s effect in inhibiting broad-spectrum biofilm. In this experiment, the biofilm from mixed cultures was formed in a 35 mm treated tissue culture dish (Hi-Media) that supports cell adherence as described previously with certain modifications (Duarte et al., 2021). The assay was divided into four sets, each having five replicates of control and treated groups as follows:

Set I (Single species biofilm of Gram-negative bacterium- *E. coli*)
Set II (Multi-strains biofilm of Gram-negative bacteria- *E. coli*) (ATCC 8739, 25922, 43888)
Set III (Multi-genera biofilm of Gram-negative bacteria- *E. coli, S. boydii, P. aeruginosa*)
Set IV (Single species biofilm of Gram-positive bacterium- *S. aureus*)

The overnight grown cultures of the above bacteria were diluted in SCDB (1×10^8^ CFU/mL) and inoculated in a tissue culture dish. For each set of the treated group, 0.1 MOI of □ ERS-1 was immediately added, while the control group was devoid of the phage treatment. Three replicates of each set were incubated for 24 h at 37 . Subsequently, the biofilms were washed with 1X-phosphate-buffered saline (PBS), pH 7.4 (Hi Media) to remove the planktonic cells, followed by quantifying three of the replicates crystal violet (CV) assay as described by (Matysik and Kline, 2019). Briefly, the biofilms were stained with 0.1% CV for 15 mins and air-dried. The dye was solubilized with acetic acid 33% (v/v), and 200 µL aliquots were transferred from each replicate into a 96-well microtiter plate (Tarsons) followed by measuring the absorbance at 595 nm with BioTek Synergy H1 microplate reader (Agilent Technologies). The microscopic examination of the biofilm was done under 100× using an OLYMPUS optical microscope U-CMAD3 T7 coupled with Lumenera Infinity 1 camera. GraphPad Prism v9.5.1 was used to compute the statistical significance of the results using RM-one-way ANOVA with multiple comparisons (FDR correction) and a p-value of 0.05.

#### 2.4.2 Time-dependent evaluation of cell viability of *E. coli* biofilm

*In-vitro* efficacy of □ ERS-1 was assessed against *E. coli* ATCC 8739 in biofilm control through inhibition and disruption assays using Film Tracer™ LIVE/DEAD® Biofilm Viability kit (Invitrogen). The experiment was carried out as described by (Mulani et al., 2022) and divided into two sets as follows:

Set I: Biofilm Inhibition (□ERS-1 treatment at 0 hr. followed by imaging the replicates at 6, 12, and 24 hrs.).
Set II: Biofilm Disruption (□ERS-1 treatment after 24 hrs. on pre-formed biofilm and imaging the replicates at 6, 12, and 24 hrs.).

For biofilm inhibition, the overnight grown host culture (*E. coli*) cells were diluted in SCDB (1×10^8^) and inoculated in a tissue culture dish, followed by the addition of □ ERS-1 at 0.1 MOI. CFU was determined after each time point of 6, 12, and 24 hours.

For biofilm disruption, the overnight grown host culture (*E. coli*) cells were diluted in SCDB (1×10^8^), inoculated in a tissue culture dish, and allowed to form biofilm for 24 hours. The planktonic cells were washed, and □ ERS-1 was added to the pre-formed biofilm at 0.1 MOI. Untreated cells of *E. coli* served as culture control. CFU was determined after each time point of 6, 12, and 24 hours. Statistical analysis of the results was performed in GraphPad Prism v9.5.1 using RM-one-way ANOVA with multiple comparisons and a p-value of 0.05.

For each time point, three replicates were used. The cells were washed with 1X-PBS (Hi Media) and stained with a mixture of SYTO^®^ 9 and propidium iodide (PI) stains from Film Tracer™ LIVE/DEAD^®^ Biofilm Viability kit following manufacturers protocol. The stained dishes were visualized with an inverted confocal laser scanning microscope (CLSM) (Leica Stellaris 5, DMi8) using a 20× objective. The fluorescence from live and dead bacteria was visualized using excitation wavelengths of 488 nm (SYTO^®^ 9) and 588 nm (PI), respectively. Additionally, to understand the effect of phage treatment on the biofilm morphology, the control and 24 h sample of disruption were imaged using Field emission scanning electron microscopy (FESEM, Nova Nano SEM 450).

### 2.5. Bacteriophage-based biocontrol for reduction of bacterial counts from wastewater

#### 2.5.1. Sample collection and details of the sampling site

Phytorid sewage treatment plant (STP) (18° 32.8246’ N, 73° 48.8338’ E) situated at CSIR-National Chemical Laboratory, India, with a capacity of 0.15 million liters per day (MLD) was chosen for sampling of untreated wastewater **(Fig. S2)**. A total of 1 L (n=3) of untreated wastewater samples were collected in a sterile bottle by grab-sampling method. The samples were brought immediately to the lab and processed for experimental setup.

#### 2.5.2. Evaluation of bacterial biocontrol by □ ERS-1 in untreated wastewater

The raw sewage (untreated wastewater) samples from Phytorid STP were divided into test and control groups. For the test group (100 mL aliquot of the raw sewage was challenged with 1 mL of high titer phage stock of □ ERS-1(10^17^ PFU/mL), while the control group was devoid of any treatment. The experiment was carried out in triplicates, and the flasks from each group were incubated at 37°C for 24 hours, followed by recording the results as CFU/mL, while the reduction in the bacterial counts was calculated in the form of log reduction and percent reduction as described previously (Bashir et al., 2022).

#### 2.5.3. Time kill assay for evaluation of bacterial biocontrol by □ ERS-1 at lab-scale

A time-kill assay was performed to harness the broad-spectrum potentials of □ ERS-1 in improving the reusability of wastewater and associated environmental health. The experimental setup comprised of two sets as follows:

Set I: Unenriched group (Test: 100 mL raw sewage +1 mL □ ERS-1), (Control: 100 mL raw sewage).
Set II: Coliform enriched group (Test: 100 mL raw sewage +10 mL Mc Conkey broth + 1 mL □ ERS-1), (Control: 100 mL raw sewage + 10 mL Mc Conkey broth).

The experiment was performed in three replicates, and aliquots from each set were drawn at 0, 6, 12, and 24 hours, followed by serial dilutions of the aliquots and plating onto selective media of Brilliant Green Bile Lactose Broth agar (BGLB-A) (Hi Media), and Eosin-methylene blue agar (EMB-A). For each time point, the bacterial count was determined as CFU/mL, and the efficacy of □ ERS-1 in reducing the bacterial counts was calculated through log reduction and (%) reduction as described by (Bashir et al., 2022). Additionally, at each time point, live-dead cell staining of the aliquots was carried out as described above and visualized under 20× objective using CLSM. Statistical analysis of the results was performed in GraphPad Prism v9.5.1 using two-way ANOVA with multiple comparisons (FDR correction) and a p-value of 0.05 for statistical significance.

## 3. Results and Discussion

### 3.1 Isolation, identification, and growth parameters of □ ERS-1

Amongst the 15 isolated phages, *Escherichia* phage (□) ERS-1 showed a broad-spectrum lytic activity against *E. coli, P. aeruginosa, S. boydii, Y. enterocolitica* and partial lysis with cloudy plaques for *S. aureus,* and therefore, was selected for detailed characterization in this study. The host range of □ ERS-1 has been tabulated **(Table S1)**.

The antimicrobial resistant profile of *E. coli* (ATCC 8739) assessed through VITEK-2 suggested its resistance toward third and fourth-generation cephalosporins (3GC, 4GC) and beta-lactam class of antibiotics, namely ceftazidime, cefepime, and Aztreonam **(Supporting information*_ E. coli_*AST281)**. The World Health Organization (WHO) declared 3GCs and 4GCs as antimicrobials of critical importance for human and animal health in 2019. However, due to an increase in the prevalence of plasmid-encoded β-lactamases in *E. coli,* members of this group have become resistant to these crucial antimicrobials (Kang et al., 2022).

The genome annotation of *E. coli* (ATCC 8739) showed the presence of β-lactamase enzymes (EC 3.5.6.2) responsible for multidrug resistance toward extended-spectrum beta-lactam antibiotics such as penicillins, cephalosporins, and aztreonam. Additionally, type 1 fimbriae are among the essential virulence factors in pathogenic *E. coli* strains and are known to promote their survival by antibiotic evasion. Besides, this strain was also found to possess resistance mechanisms (enzymes and efflux pumps) for multiple antibiotics **(Table S2)**. Therefore, the isolation of phage for improved management of *E. coli* and control of resistance transfer for 3, 4 GCs, and multidrugs highlights the importance of this study.

The plaque morphology of □ ERS-1 with this host comprised clear plaques encircled in a translucent halo **(Fig 1A)**. This type of plaque morphology could be attributed to the occurrence of polysaccharide depolymerases in either the structural proteins or tail fibers of □ ERS-1. These enzymes are non-lytic and are known to cleave the extracellular polysaccharides of bacteria, reducing their virulence (Rice et al., 2021). This preliminary observation suggested that□ERS-1 could have potentials antibiofilm activity.

**Figure 1:**
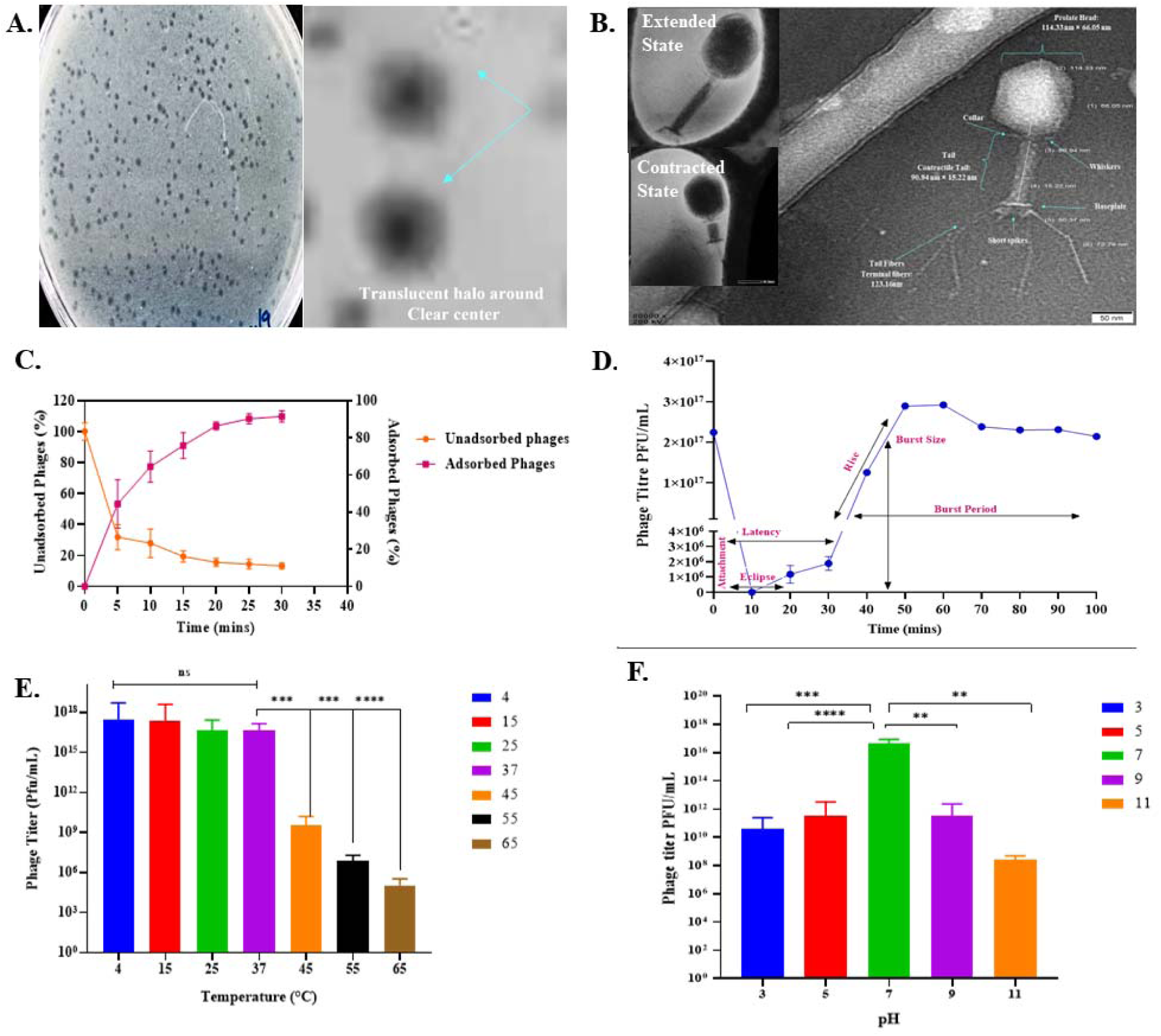
Morphology and growth parameters of □ ERS-1. **(A)** Plaque morphology. The blue arrows point to the zoomed view of plaques, showing a clear center surrounded by a translucent halo. **(B)** HR-TEM micrographs of □ ERS-1 at 80000 ×, and scalebar of 50 nm. The inset is STEM images of □ ERS-1 in extended and contracted states. **(C)** Graphical representation of the percentage of adsorbed and unadsorbed phage particles of □ ERS-1 as a function of time. **(D)** One-step growth curve of □ ERS-1 with the phases of its lifecycle marked (pink). **(E)** Illustration of the effect of temperature on the stability of □ ERS-1. **(F)** Effect of various pH ranges on the stability of □ ERS-1. Data in the graph panels from C-F represent means of the experimental replicates, standard deviations (SD), and statistical significance of the data (*, **, ***, ****) using RM-one-way ANOVA. The acronym ‘ns’ implies no significant difference between the test groups.

A detailed morphological characterization of the purified high titer stock of □ ERS-1 was carried out with HR-TEM and STEM, which revealed typical “T4-like” features of myovirus. The morphological architecture of □ ERS-1 comprised of a prolate head (length, 114 ± 5 nm; width, 66□±□3 nm) showing an icosahedral symmetry, a collar with whiskers, a contractile tail (length, 90□±□10 nm; width, 15□±□2 nm), a small baseplate with short spikes, and six long terminal (tail) fibers **(Fig 1B)**. Based on the above morphological features, □ ERS-1 was classified into the class *Caudoviricetes* and family *Straboviridae,* as suggested by the recent taxonomic update of ICTV (Turner et al., 2023).

The titer of □ ERS-1 was determined as 4.8×10^17^ PFU/mL. An MOI of five or greater indicates the host cell being infected by more than one phage particle, thereby portraying the virulent ability of phage to set up a productive infection within the susceptible host (Abedon, 2023). The MOI for □ ERS-1 was calculated as 8.8, which could achieve a bacterial clearance of 99.98%, with a probability of infected cells being P_0_=0.998. This indicates the virulent nature of □ ERS-1 and its capability of mounting a productive infection in *E. coli* (ATCC 8739) cells.

The binding of the phage to specific receptors on the bacterial cell surface initiates bacteriophage infection of the susceptible host. This event is termed phage adsorption, an essential parameter in deciding the application of a bacteriophage (Ge et al., 2020). The percentage of adsorbed fraction of □ ERS-1 particles was 50% up to 5 mins, which increased to >90% by 20 mins. Additionally, the percentage of free phages in the unadsorbed fraction decreased rapidly at 5 mins (∼70%), and only ∼10 % of free phages were observed from 20-30 mins **(Fig 1C)**. Details of the adsorption rate have been summarised **(Table S3)**.

The fundamental nature of replication of □ ERS-1 in *E. coli* (ATCC 8739) was determined with one step growth experiment. This enables us to understand the duration of the different phases in the lifecycle and the burst size (yield of the viral cycle) (Adams and Wassermann, 1956). Phage □ ERS-1 showed a latent period of 20 mins and a burst size of 45 (±5) phage particles per infected cell of *E. coli* (host) **(Fig 1D)**.

Various external factors are known to affect phage persistence. However, according to Ackermann, tailed phages tend to be most stable under adverse external factors. The temperature is a crucial factor in determining the survival and infection cycle of the phage. This factor is also essential for phages’ short-term and long-term storage to retain their activity (Jończyk et al., 2011). The temperature stability of □ ERS-1 was determined from 4□ to 65□. Interestingly, it was observed that cold conditions of 4 and 15□ were found to be optimal temperatures as the phage titer increased from 10^17^ PFU/mL to 10^20^ PFU/mL. This could be because the phage was isolated from the Ganges River’s upper stretch (Harshil location), which has a relatively cold temperature. An increase in the temperature from 37□ to 45□ resulted in a significant loss in the phage titer (ANOVA, p<0.001). Additionally, a significant decline was observed in the phage titer at 55□ and 65□ (ANOVA, p<0.001) **(Fig 1E)**.

Another essential factor that limits the stability and activity of phages is the pH of the environment(Jończyk et al., 2011). The pH stability of □ ERS-1 was determined from pH 3 to 11. From the phage titration, pH 7 was the optimum pH for sustaining phage stability. On either side of the pH scale, it was observed that the phage titer reduced from 10^17^ PFU/mL to 10^10^ PFU/mL (pH 3) and 10^12^ PFU/mL (pH 5, pH 9), respectively **(Fig 1F)**. However, a phage titer of 10^9^ PFU/mL is considered a high titer (Bonilla and Barr, 2018). This suggests that □ ERS-1 continues to have an infectious nature for *E. coli* even at a diverse pH range. Additionally, in the environment, especially in wastewater, most microbes thrive at pH 6-9; therefore, most of the biological treatment of wastewater occurs at this pH range (Bouchaala et al., 2021). The ability of □ ERS-1 to mount infection to its target host at various ranges of temperature and pH substantiates its candidacy for use in wastewater treatment.

### 3.2 Genome analysis, annotation, and proteome of □ ERS-1

The genome length of □ ERS-1 consists of 175,242 bp and 60.9% of G+C content. Analysis of ORFs revealed the presence of 796 ORFs, mainly comprising hypothetical proteins followed by phage proteins involved in defining phage structure, DNA replication, repair, packing, and lysis of the host. The details and position of various proteins in the genome of □ ERS-1 have been illustrated in **Fig 2A**. Phage lysins are considered highly efficient and evolved molecules that can digest the peptidoglycan in the cell wall of bacteria, thereby facilitating the release of viral progeny (Fischetti, 2005). From genome annotation using RAST and Prokka v1314.6 tool, it was evident that lysin was the key enzyme responsible for the host lysis by □ ERS-1 .

**Figure 2:**
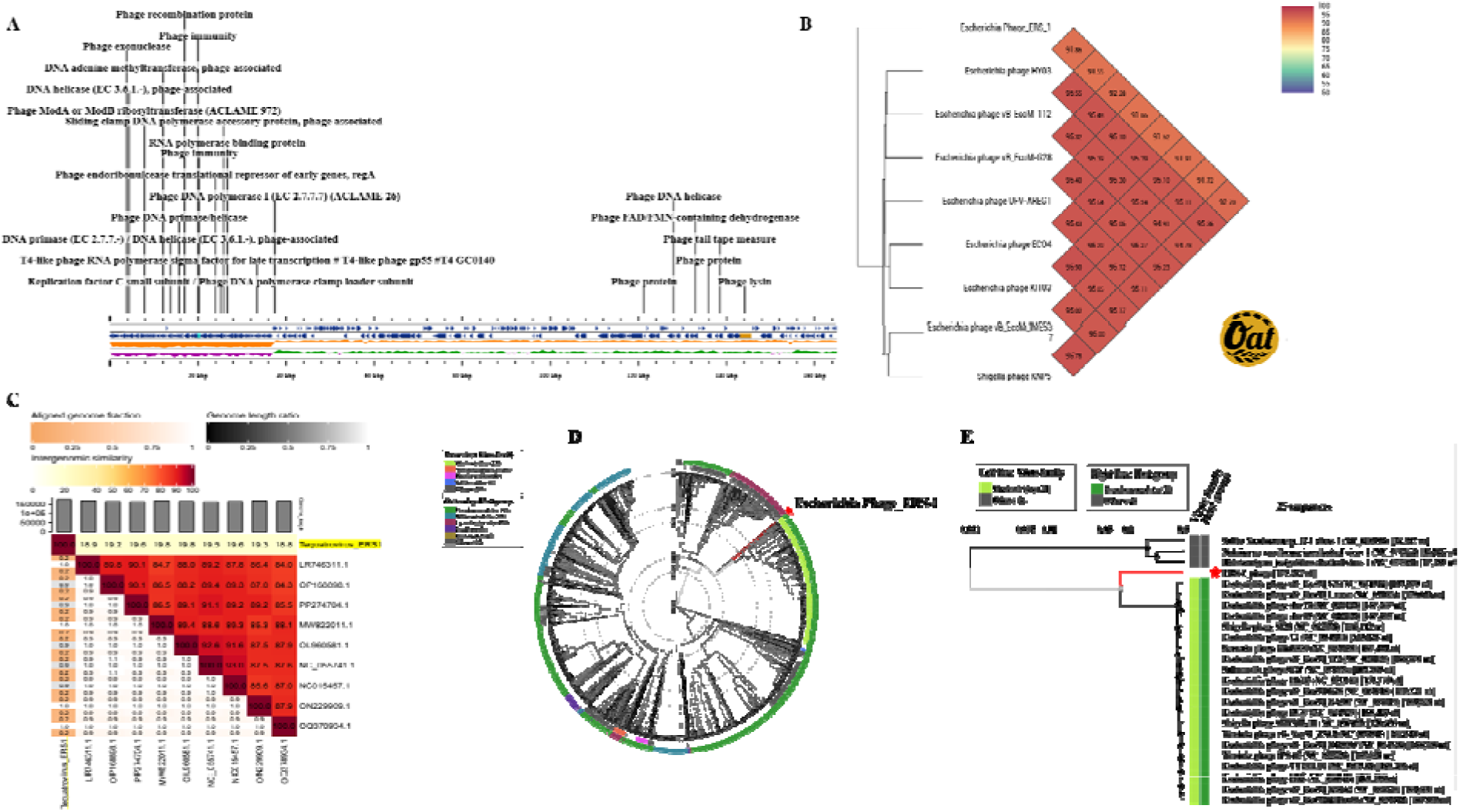
Genome details and amino-acid-based comparative genomic and phylogenetic analysis of □ ERS-1. **(A)** Linear genome map of □ ERS-1 created with Proksee viewer. The outer blue arrows are the RAST annotated protein-coding genes on the positive and negative strands of the DNA. The inner orange skew displays the total GC content, followed by GC skew information in the positive (green) and negative (purple) strands. **(B)** Heatmap of OrthoANI between □ ERS-1 and closely related *Tequatroviruses*, computed with OAT software. The color scale (top right) contains values defining the similarity percentage between the genomes. The numerical values of intergenomic genetic distance (from OAT software) are marked on the branches. **(C)** Heatmap showing alignment indicators (left half) and intergenomic similarity values (right-half) generated through VIRDIC. **(D)** A circular phylogenetic tree generated by VipTree for all 3036 reference genomes of phages related to □ ERS-1. The genome distance matrix of the phage under consideration and its relatives are calculated by BIONJ (an algorithm based on the distance for phylogeny reconstruction (Gascuel, 1997). The midpoint of the tree marks the root. The outer-colored rings (host taxa) and the inner-colored rings (viral families). **(E)** Based on the SG scores, a rectangular phylogenetic tree of □ ERS-1 with its 21 closely related and 3 distantly related phage genomes. The red stars represent Escherichia phage □ ERS-1.

Further, from the orthoANI comparison of the genome of □ ERS-1 and closely related *Tequatrovirus* members, it was observed that □ ERS-1 has an ANI of 94.2% with Escherichia phage LH2, indicating the close relatedness of its genome **(Fig 2B)**. Additionally, the intergenomic similarities analysis with VIRDIC tool revealed only 19% similarity of this phage with its closest relatives, indicating distant relatedness of the genome of phage □ ERS-1 **(Fig 2C)**. However, VIRIDIC is not able to capture similarity relationships between the related viruses which have regions of similarity of less than 65%, a limitation inherent to BLASTN. Therefore, protein-based analysis is recommended to clarify the phylogenetic relationships. Analysis of amino-acid sequences in the whole genome of □ ERS-1 with all the available reference phages, and closely related phages was performed using VipTree. A total of 3036 reference sequences of the phage genomes were used to construct the phylogenetic tree **(Fig 2D)**. The preliminary observation of the circular phylogenetic tree of □ ERS-1 was consistent with morphological features of ‘T4 like myovirus’. Further, depending on the S_G_ scores of tblastx, a subset of 21 closely related phages and three distantly related phages were chosen to construct a rectangular phylogenetic tree **(Fig 2E)**. The genome of □ ERS-1 clustered with some *Escherichia* phages, followed by *Shigella* phages, all belonging to *Tequatrovirus,* a genus of the *Straboviridae* family, according to ICTV. As per ICTV, the isolated phage(s) are classified as a member of the same genus if their identities of nucleotide sequences are >70% and a distinct species if ANI is ≤95% (Bin Jang et al., 2019). The above nucleotide and amino-acid-based analysis of □ ERS-1 suggested that □ ERS-1 could be classified as a new member within the *Straboviridae* family and *Tequatrovirus* genus. The genome of this phage has been submitted to NCBI and has been assigned accession number PP337211.

The members of *Tequatrovirus* are known to infect *Escherichia, Enterobacteria, Shigella, Salmonella, Yersinia, Aeromonas, Burkholderia, Stenotrophomonas, Prochlorococcus, Synechococcus, Citrobacter* and *Staphylococcus* (https://www.ncbi.nlm.nih.gov/Taxonomy/Browser/wwwtax.cgi?id=10663). However, it is noteworthy that there are no reports of *Tequatrovirus* infecting *P. aeruginosa* in the NCBI database. Hence, to our knowledge, this could be the first report detailing a complete genome of a *Tequatrovirus* member capable of infecting *P. aeruginosa* ATCC 9027.

### 3.3 Anti-Biofilm Potential of □ ERS-1

#### A. Single, Multi-strain, Mixed-Genera Biofilm Inhibition Spectra

Several reports have suggested the occurrence of single species biofilm and mixed or multi-species/ mixed biofilms in nature. The most frequently encountered bacteria in formation of biofilm in environment, medicine, food industry, agriculture and animal husbandry include members of *Escherichia, Shigella, Salmonella, Pseudomonas, Acinetobacter, Klebsiella, Enterococcus, Streptococcus,* and *Staphylococcus* (Burmølle et al., 2014; Denissen et al., 2022; Mouiche et al., 2019; Sharma et al., 2023; Toushik et al., 2022). Given this, we evaluated the potentials of □ ERS-1 for inhibition of biofilm by single species, multiple strains, and multiple genera, using the information of its host range obtained through spot assay. Results of the CV assay indicated that □ ERS-1 could inhibit and reduce 96.17% of the biofilm mass produced by *E. coli* (ATCC 8739) **(Fig 3A, E)**.

**Figure 3:**
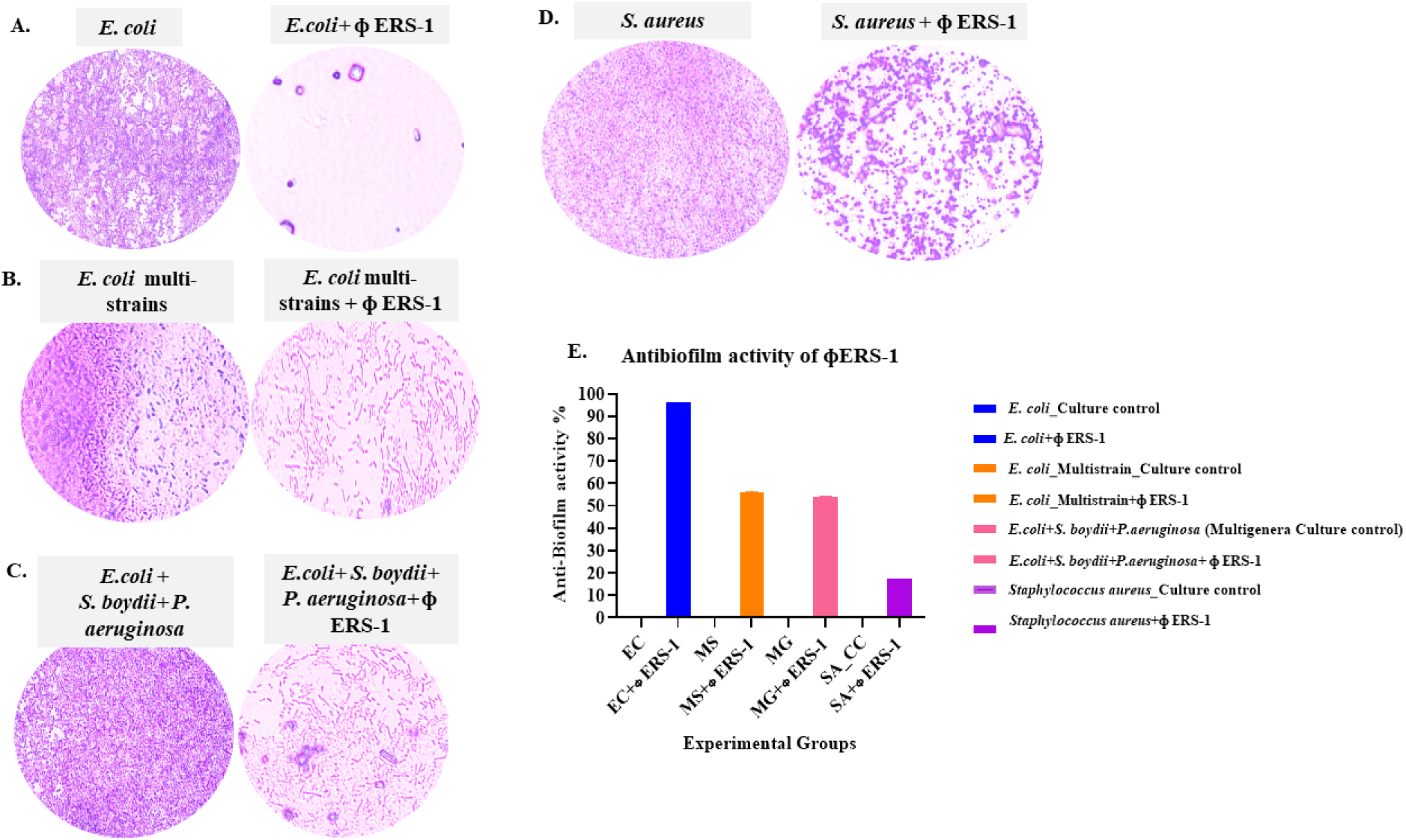
Effects of □ ERS-1 in biofilm inhibition by single species, multi-strains, and multiple genera of bacteria with CV assay. Figure 3A-D are CV-stained microscopic images of biofilm, observed under an oil immersion lens (100×) after the control and treated groups were incubated for 24 hours. **(A)** Single species biofilm of *E. coli* culture control (left), *E. coli* treated with □ ERS-1 (right). **(B)** Multiple strains of *E. coli* mixed to form biofilm-culture control (left), multi-strain group treated with □ ERS-1 (right). **(C)** Biofilm formation by multiple genera – culture control (left), multiple genera group treated with □ ERS-1 . **(D)** Biofilm formation by *S. aureus-* culture control (left), and □ ERS-1 treated (right). **(E)** Quantification of biofilm formation with CV assay, followed by absorbance at 590 nm.

Interestingly, it was observed that the addition of □ ERS-1 to multiple strains of *E. coli* (ATCC 8739, 25922, 43888) **(Fig 3B, E)** and multiple genera (*P. aeruginosa* (ATCC 9027), *S. boydii* (ATCC 9207), and *E. coli* (ATCC 8739) **(Fig 3 C, E)** inhibited the formation of biofilm by 56.2% and 54% respectively. This is an important observation, as bacterial biofilms are more resistant to external factors and treatment by antimicrobial and chemical agents than their planktonic counterparts. The ability of □ ERS-1 to render the cells in their planktonic form could be attributed to the presence of polysaccharide depolymerases in either free or bound state, which is defined by the semi-transparent halo around phage plaques as evident in **(Fig 1A)**. The other plausible mechanism behind the broad-spectrum antibacterial and antibiofilm nature of □ ERS-1 could be the production of certain quorum-quenching enzymes or lipases. Quorum-quenching enzymes in phages are responsible for inhibiting quorum sensing by mixed species/genera biofilms composed of *P. aeruginosa* and *E. coli* (Liu et al., 2022; Pei and Lamas-Samanamud, 2014). On the other hand, lipases disperse biofilms by hydrolyzing lipids in the bacterial cell membrane (Azeredo et al., 2021). However, there is limited information about these enzymes. Therefore, to have a holistic understanding of the mechanism of bacterial lysis and broad host specificity, an in-depth study of phage-derived proteins of □ ERS-1 is necessary.

#### B. Time-dependent evaluation of *E. coli* biofilm inhibition and disruption

*E. coli* is a prominent fecal indicator in the waterways. The biofilm formation ability of *E. coli* on the surface of water treatment pipes and filters poses a significant challenge for treating and preventing this bacteria during disinfection and recycling practices of water and wastewater (Qian et al., 2022; Steven et al., 2022). From the genome annotation study, the biofilm composition of the *E. coli* host in this study was linear homopolymer poly-beta-1,6-N-acetyl-D-glucosamine (beta-1,6-GlcNAc; PGA) **(Table S2)**. The pgaABCD operon’s gene products are necessary for forming and maintaining biofilm structural stability in several enteric pathogenic *E. coli* (Itoh et al., 2008). Therefore, in this study, we evaluated the potentials of □ ERS-1 in the prevention (inhibition) and dispersal (disruption) of biofilm by *E. coli* ATCC 8739 in a time-dependent manner.

The CLSM analysis of □ ERS-1 treated *E. coli* showed red fluorescence at 6, 12, and 24 hours, indicating the dead biomass of *E. coli* cells **(Fig 4 B-D)**, as compared to the control (devoid of □ ERS-1 treatment) **(Fig 4A)**. The presence of live biomass was ascertained through CFU counts of viable cells. Results indicated a significant inhibitory effect on biofilm formation at 6 hrs., which continued up to 24 hours (ANOVA, p=<0.0001) **(Fig 4D)**. Previous studies have demonstrated the antibiofilm effect of *Tequatrovirus* against *E. coli* and *Salmonella,* with bacterial reduction from 2-5 logs within 6-24 hours (Liao et al., 2022; Zhou et al., 2022). However, in our study, for the first time we report the potential of □ ERS-1 in reducing the abundance of *E. coli* cells from 6.14 log_10_ CFU/mL at 6 hours to 8.22 log_10_ CFU/mL at 24 hours, with a ∼100% reduction in the total viable counts. This data suggests that, ERS-1 poses an inhibitory effect that could be used for biological control of surface colonization by *E. coli* cells either through inhibition of their initial attachment or impediment of bacterial establishment on various surfaces.

**Figure 4:**
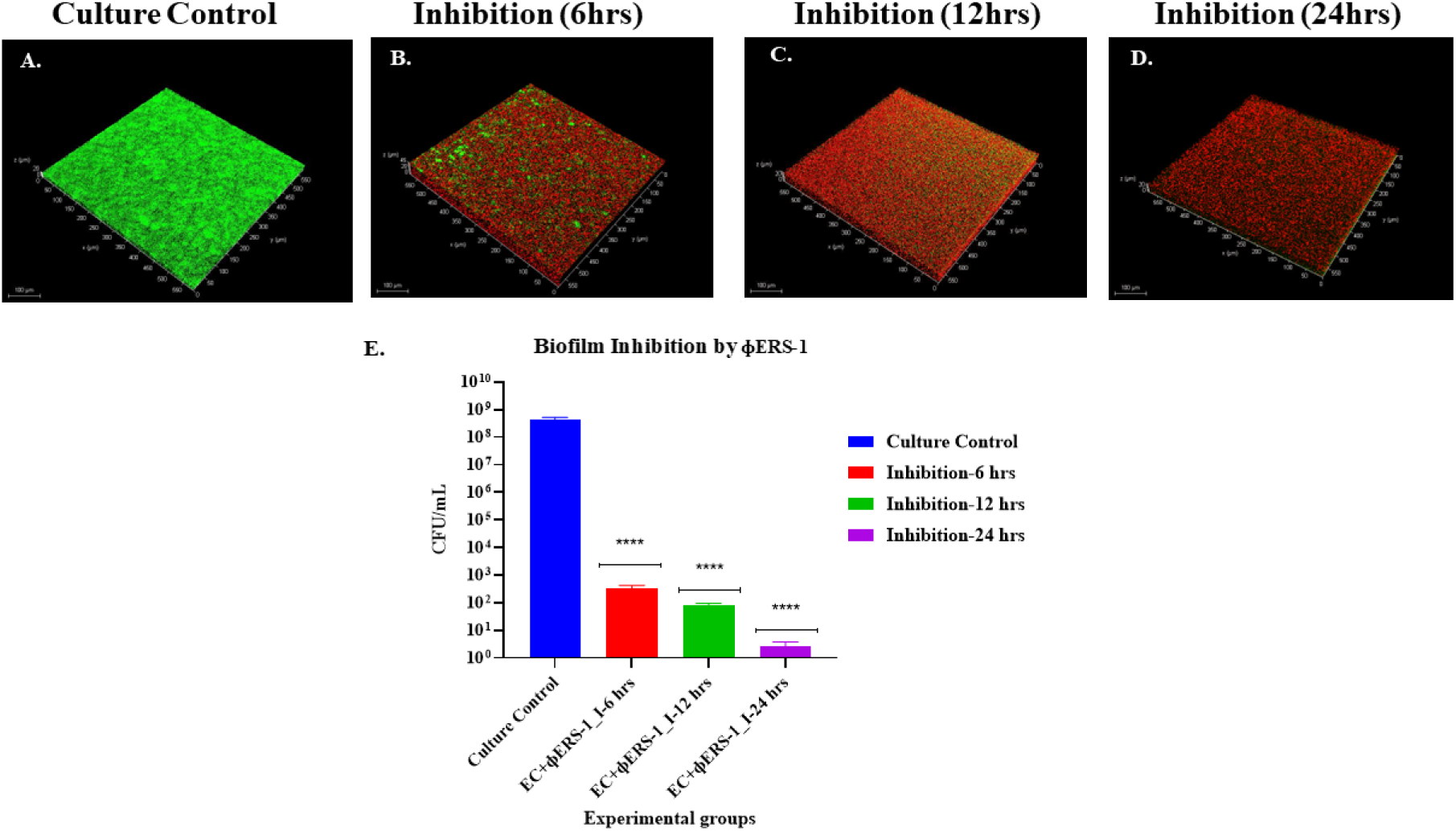
Inhibitory effect of □ ERS-1 on biofilm formation by *E. coli.* Samples were observed under a Leica Stellaris 5, DMi8 microscope with a 20× objective. The red fluorescence indicates dead *E. coli* cells, while the green fluorescence indicates viable cells at different time intervals. **(A)** CLSM micrograph of *E. coli* cells with no treatment, capable of forming biofilm after 24 hours. **(B)** CLSM micrograph of □ ERS-1 treated group imaged after 6 hours. **(C)** CLSM micrograph of □ ERS-1 treated group imaged after 12 hours. **(D)** CLSM micrograph of □ ERS-1 treated group imaged after 24 hours. **(E)** Quantification of the viable cells as CFU/mL of the control (non-treated) and □ ERS-1 treated sets.

In the natural environments, most of the biofilms are in their mature state, and therefore, to detach the biofilm, disruption of EPS polymers followed by planktonic cell dispersal is essential (Shrestha et al., 2022). Capsular polysaccharides as major virulence factors have been reported for *A. baumannii* (Wang et al., 2020)*, K. pneumoniae* (García et al., 2019), and *E. coli* (Guo et al., 2017) in biofilm formation. Thus, the antibiofilm effect of □ ERS-1 in disintegrating matured or pre-formed biofilms was evaluated with CLSM and counts of viable cells using CFU/mL.

It was observed that the number of dead/ non-viable cells increased in the phage-treated group for 24 hours **(Fig 5A-D)**. This indicates a gradual disruption of *E. coli* biofilm by □ ERS-1. The total viable counts at each time point revealed that □ ERS-1 was capable of disrupting the pre-formed biofilm layer of *E. coli* with a reduction of 2.4 log_10_ CFU/mL at 6 hours, followed by 2.70 log_10_ CFU/mL at 12 hours, and 3.88 log_10_ CFU/mL at 24 hours. These values indicate that □ ERS-1 can disrupt biofilm with a percent reduction of viable cells up to 99% with 6 hours of treatment. After 24 hours, the percent reduction of *E. coli* cells observed in the biofilm was 99.98% (ANOVA, p=0.0022). **(Fig 5 K)**.

**Figure 5:**
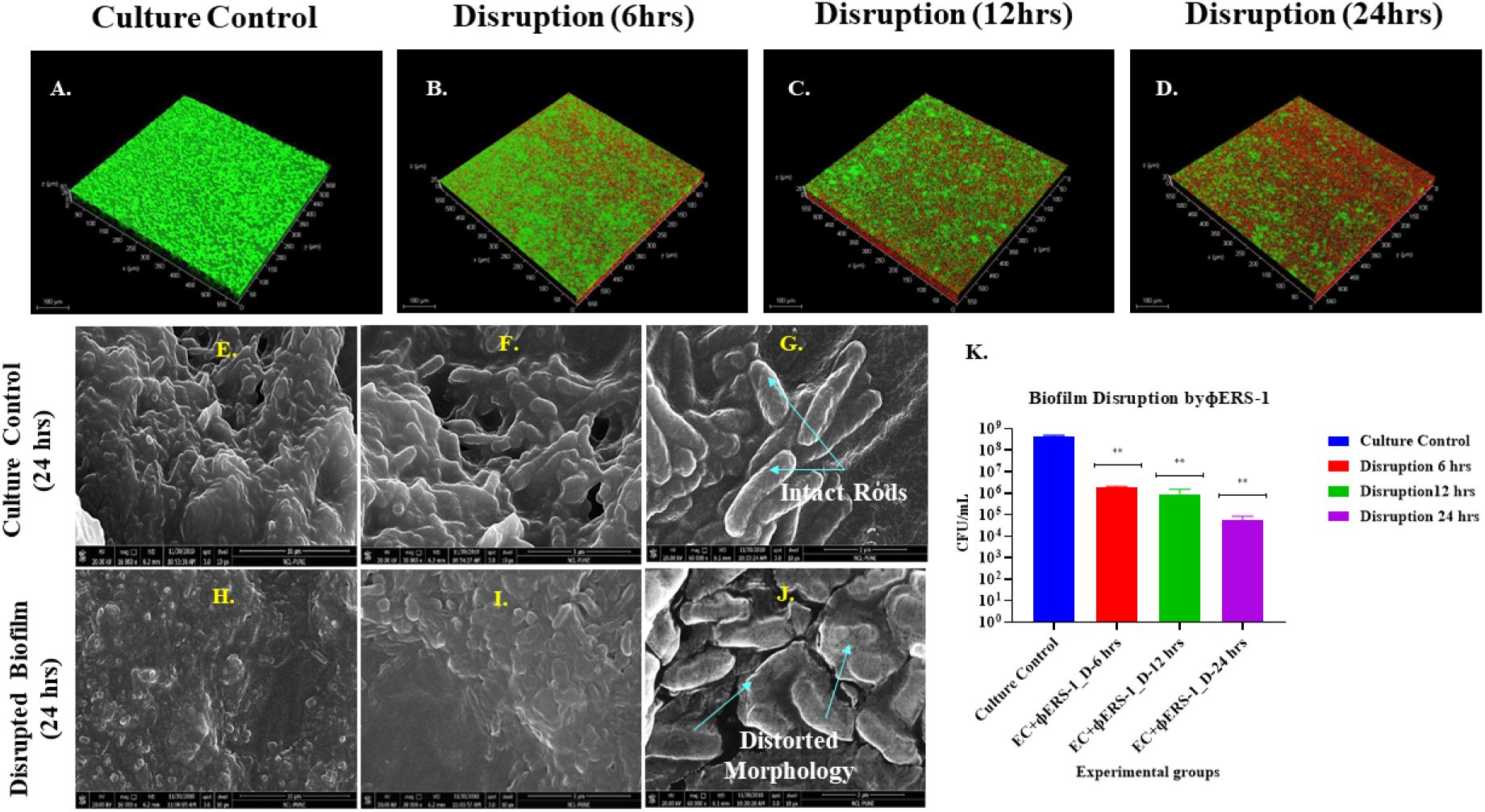
Effect of □ ERS-1 on pre-formed/ matured biofilm of *E. coli* (ATCC 8739). The control and treatment sets visualized through CLSM with 20 × objective at specified time intervals are represented in the panels as **(A)** CLSM micrograph of culture control with pre-formed biofilm of *E. coli* cells devoid of any treatment. **(B-D)** CLSM micrograph of □ ERS-1 treated pre-formed biofilm imaged after 6 hours, 12 hours **(C)**, and 24 hours **(D)**. **(E-G)** FE-SEM images of control biofilm at different magnifications 16000× **(E)**, 30000× **(F)**, and 60000× **(G)**, showing intact rods immersed in the EPS matrix of the biofilm. **(H-J)** FE-SEM images of □ ERS-1 treated biofilm at different magnifications 16000× **(H)**, 30000× **(I)**, and 60000× **(J)**, showing disrupted EPS matrix and distorted morphology of compromised cells in the biofilm. **(K)** Quantification of the viable cells in disrupted biofilm as CFU/mL of the control (non-treated) and □ ERS-1 treated sets.

Additionally, the FESEM imaging of the control and □ ERS-1 treated biofilm after 24 hours confirmed disruption of the pre-formed biofilms. The cells of the control set showed a well-defined, rod-shaped morphology embedded in a matrix of EPS **(Fig 5 E-G),** while the □ ERS-1 (24 hrs.) treated set showed a distorted morphology within the biofilm **(Fig 5 H-J)**. The planktonic bacteria within biofilms are highly resistant to the action of antibiotics, disinfectants, and disruption by physical or chemical ways (Kovacs et al., 2023). However, the experimental evidence in this study suggests the efficacy of □ ERS-1 in biofilm disruption and its bacteriolytic effects against the planktonic cells, demonstrating its potential applications.

### 3.4 Bacteriophage-based reduction of bacterial counts from wastewater

The disinfection of the WW to make it safe is crucial to protecting the health of the environment, humans, and animals (Regulation EU 2020/741). Failure to do so could be responsible for different diseases because of various pathogens. According to this regulation, the reclaimed water is monitored based on various pathogen indicators. For bacteria: *E. coli;* for pathogenic viruses: coliphages; and protozoa: the spores of *Clostridium perfringens* or sulphate-reducing bacteria (Hernández-Crespo et al., 2022; Kadlec and Wallace, 2008). Therefore, considering the broad-spectrum host range, bacteriolytic, and antibiofilm potentials of □ ERS-1, we focused its application on reducing the bacterial load from untreated wastewater/ raw sewage.

The initial screening results showed a significant reduction of the bacterial load of 2.27 log_10_ CFU/mL in the □ ERS-1 treated group of the raw sewage after 24 hours (p<0.001) **(Fig 6 A, B)**. Further, a time-kill assay was performed to evaluate the magnitude of reduction in the viable coliform load in control and □ ERS-1-treated groups. The rationale behind plating the raw sewage and coliform enriched (in McConkey broth) on BGLB agar and EMB agar was differentiation and specific enumeration of the coliform bacterial load. Results after 24 -48 hours of incubation revealed a significantly lower number of viable bacteria in the □ ERS-1 treated group in both raw sewage (ANOVA, p<0.01) and coliform enriched group (ANOVA, p<0.001). Interestingly, in the set of □ ERS-1 treated raw sewage, after 24 hours, a 2-2.4 log_10_ reduction was observed compared to the control group. Notably, in the coliform enriched set of raw sewage, 4.2 log_10_ CFU/mL reduction was observed in the □ ERS-1 treated group **(Fig 6 C-E)**. Additionally, the CLSM imaging of aliquots also revealed an increase in the dead cells (red) over time in the □ ERS-1 treated group of unenriched and coliform-enriched sewage **(Fig 6 F)** compared to the control group. This indicates that □ ERS-1 could be used as a green approach to reduce the coliform counts.

**Figure 6:**
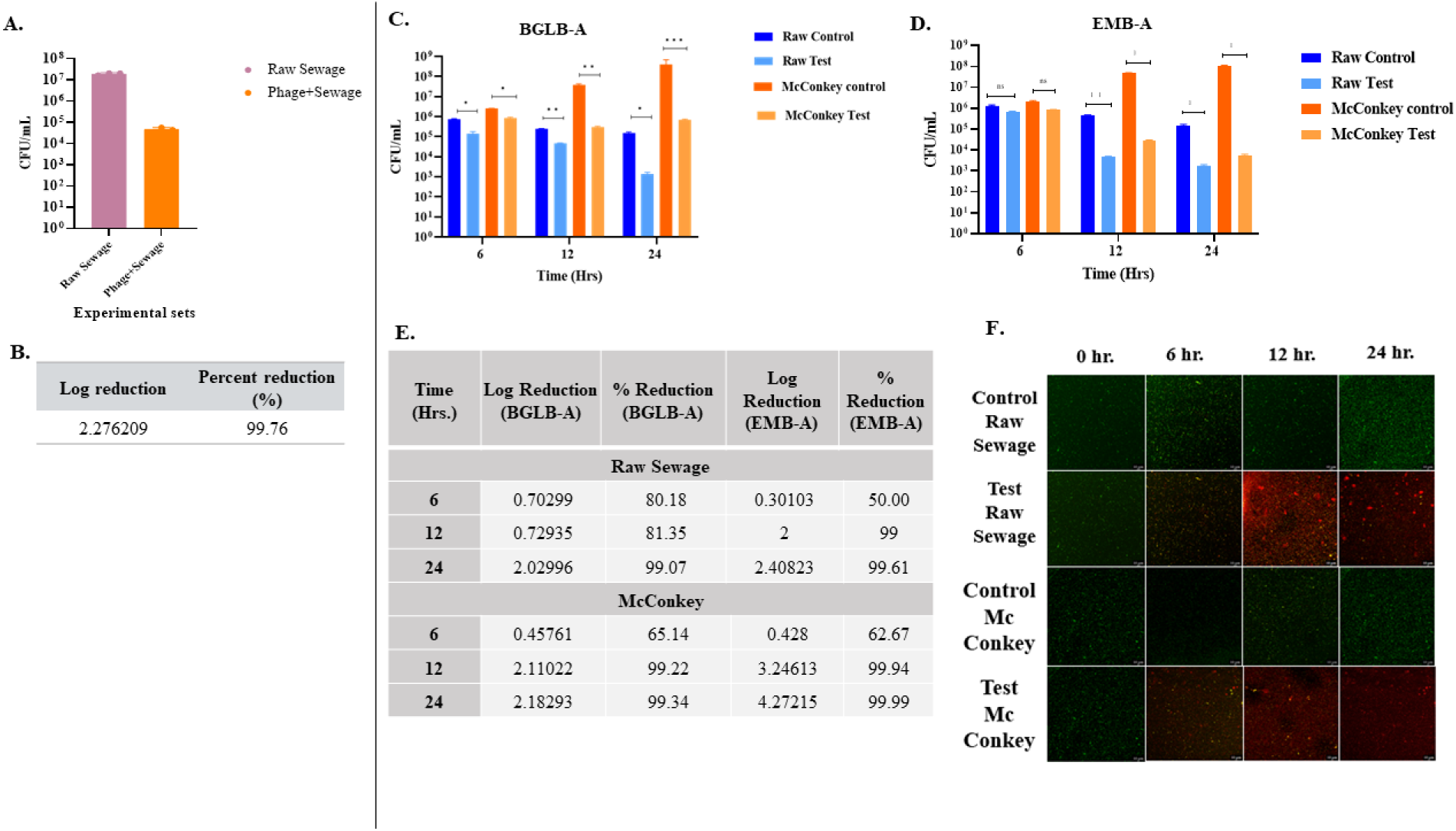
Evaluation of □ ERS-1 in reduction of coliform counts. **(A)** Graphical representation of quantification of the viable cells in raw sewage as CFU/mL of the control (non-treated) and □ ERS-1 treated sets. **(B)** Log reduction and percentage reduction values indicate □ ERS-1 efficacy in reducing bacterial load in raw sewage after 24 hours. **(C, D)** Graphical representation of quantification of the viable cells in raw and coliform enriched sewage as CFU/mL of the control (non-treated) and □ ERS-1 treated sets on BGLB-agar **(C)**, and EMB agar **(D)**, as a function of time. **(E)** Log reduction and percentage reduction values indicate the time-dependent efficacy of □ ERS-1 in reducing bacterial load in raw and coliform-enriched (McConkey broth) sewage. **(F)** CLSM micrographs showing live (green) and dead (red) bacterial cells in the raw (unenriched) sewage and coliform-enriched (McConkey) sewage at different time intervals.

There are few reports describing the use of phages in wastewater treatment to reduce the bacterial load and the use of phage cocktail to eliminate waterborne pathogenic bacteria (Grami et al., 2022; Jassim et al., 2016; Periasamy and Sundaram, 2013). However, to the best of our knowledge, this is the first study to highlight the use of single polyvalent, non-engineered (naturally occurring) phage □ ERS-1 to reduce the coliform load in wastewater. Also, this is the very first kind of study detailing an in-depth characterization, antibacterial, and antibiofilm potentials, together with lab-scale evaluation of reduction in coliform counts in raw sewage by a novel phage □ ERS-1 isolated from the untapped location of the Ganges River. However, a detailed understanding of the optimized phage dose and its effect on physicochemical and microbiological parameters before and after □ ERS-1 treatment would be necessary for its on-site application. Additionally, to overcome resistance by bacteria in the long run, combining various phages with □ ERS-1 or using phage-derived enzymes from □ ERS-1 would benefit its application under one health canopy, promoting human, animal, and environmental health.

## 4. Conclusion

This study demonstrates the isolation and characterization of a novel phage □ ERS-1 from an untapped location of the Ganges River. The strength of our research lies in the novelty of phage and its bacteriolytic and antibiofilm potentials in reducing the bacterial load of environmentally relevant waterborne pathogens in their planktonic cells and biofilm form. The concept of controlling bacteria using phages and their enzymes without the aid of chemical or UV disinfection is a challenging one. However, the data also shows the efficacy of the newly isolated, high titer □ ERS-1 in controlling the bacterial (coliform) load in the sewage water and possibly improving the water quality. Our approach of using polyvalent phage as a green alternative for bacterial biocontrol of sewage water highlights the importance of the potential applications of such broad-spectrum bacteriophages to improve the quality of the effluent and disposal of sludge in the environment. This approach can significantly impact the delivery of sustainable development goals 6: clean water and sanitation, 3: good health and well-being, and 11 sustainable cities and communities. Moreover, using lytic phages to improve wastewater quality would substantially reduce the load on other treatment methods, thereby contributing to low energy consumption levels, reduced use of harmful chemicals, and improved access to water reuse with reduced biological contamination, promoting a circular economy.

## 5. Credit authorship contribution statement

**RS:** Conceptualization, visualization, experimentation, sampling, writing-review & editing, and Data analysis. **KK:** Project monitoring, Sample collection, editing, **MSD:** Conceptualization, Supervision, review, and editing.

## 7. Acknowledgments

Authors are thankful to the National Mission for Clean Ganga (NMCG), Government of India, New Delhi, India, for the project (GKC-01/2016-17, 212, NMCG-Research), Directors of CSIR-NCL, and CSIR-NEERI for infrastructure and support. RS acknowledges Mr. Manan Shah for his help in sampling. RS is grateful to Mr. Tushar Kolhe and Mr. Chetan from Central Analytical Facility, CSIR-NCL, for their help in HR-TEM and FE-SEM. RS is thankful to HRDG-CSIR and NMCG, New Delhi, for fellowship and AcSIR, New Delhi, for the academic support. The manuscript has been checked for plagiarism using iThenticate software with an institutional license.

