## Supplementary material for "Characterization of a Novel *Tequatrovirus* Phage from Pristine Stretch of The Ganges River, India, in Reducing Bacterial Load from Sewage Water": Ecoli_AST281_R-1.pdf

bioMerieux Customer: CSIR-NCL
System #: VK2C14748

### Laboratory Report

Printed Nov 26, 2020 12:15 IST
Printed by: ss.jamdade
Report Version: 3 of 3

Isolate Group: Ecoli_AST281_R-1

Card Type: AST-N281 Testing Instrument: 00001766B68C (14748)

Organism Quantity:
Selected Organism: Escherichia coli

| Comments: |
|---|

| Identification Information | | | | |
|---|---|---|---|---|
| Selected Organism | Escherichia coli | | | |
| | Entered: | Nov 15, 2020 16:58 IST | By: | ss.jamdade |

**Analysis Messages:**

The following antibiotic(s) are suppressed from analysis:
Ticarcillin/Clavulanic Acid,

| Susceptibility Information | Card: | AST-N281 | Lot Number: | 7011215403 | Expires: | Mar 25, 2021 12:00 IST |
|---|---|---|---|---|---|---|
| | Completed: | Nov 16, 2020 07:29 IST | Status: | Final | Analysis Time: | 14.50 hours |

| Antimicrobial | MIC | Interpretation | Antimicrobial | MIC | Interpretation |
|---|---|---|---|---|---|
| Ticarcillin/Clavulanic Acid | | | Amikacin | <= 2 | S |
| Piperacillin/Tazobactam | <= 4 | S | Gentamicin | <= 1 | S |
| Ceftazidime | <= 1 | *R | Ciprofloxacin | 0.5 | S |
| Cefoperazone/Sulbactam | <= 8 | S | Levofloxacin | <= 0.12 | S |
| Cefepime | <= 1 | *R | Minocycline | 4 | S |
| Aztreonam | <= 1 | *R | Tigecycline | <= 0.5 | S |
| Doripenem | <= 0.12 | S | Colistin | <= 0.5 | S |
| Imipenem | <= 0.25 | S | Trimethoprim/Sulfamethoxazole | <= 20 | S |
| Meropenem | <= 0.25 | S | | | |

+= Deduced drug *= AES modified **= User modified

| AES Findings: | Last Modified: | Mar 19, 2017 12:30 IST | Parameter Set: | Copy of Global CLSI-based+Natural Resistance |
|---|---|---|---|---|
| Confidence Level: | Consistent | | | |

Installed VITEK 2 Systems Version: 07.01
MIC Interpretation Guideline: Copy of Global CLSI-based
Therapeutic Interpretation Guideline: NATURAL RESISTANCE
AES Parameter Set Name: Copy of Global CLSI-based+Natural Resistance
AES Parameter Last Modified: Mar 19, 2017 12:30 IST

bioMerieux Customer: CSIR-NCL
System #: VK2C14748

### Laboratory Report

Printed Nov 26, 2020 12:15 IST
Printed by: ss.jamdade
Report Version: 3 of 3

Isolate Group: Ecoli_AST281_R-1

Card Type: AST-N281 Testing Instrument: 00001766B68C (14748)

---

Organism Quantity:
Selected Organism: Escherichia coli

| Action | Name (User ID) | Date/Time | Comment |
|---|---|---|---|
| **Reviewed by:** | **Sayali Jamdade (ss.jamdade)** | **Nov 21, 2020 16:30 IST** | |
| **Approved by:** | **Syed g Dastager (SyedDastager)** | **Nov 21, 2020 16:35 IST** | |

---

Installed VITEK 2 Systems Version: 07.01
MIC Interpretation Guideline: Copy of Global CLSI-based
Therapeutic Interpretation Guideline: NATURAL RESISTANCE
AES Parameter Set Name: Copy of Global CLSI-based+Natural Resistance
AES Parameter Last Modified: Mar 19, 2017 12:30 IST
