## Supplementary figures and images for "Characterization of a Novel *Tequatrovirus* Phage from Pristine Stretch of The Ganges River, India, in Reducing Bacterial Load from Sewage Water"

### Fig S1.tif

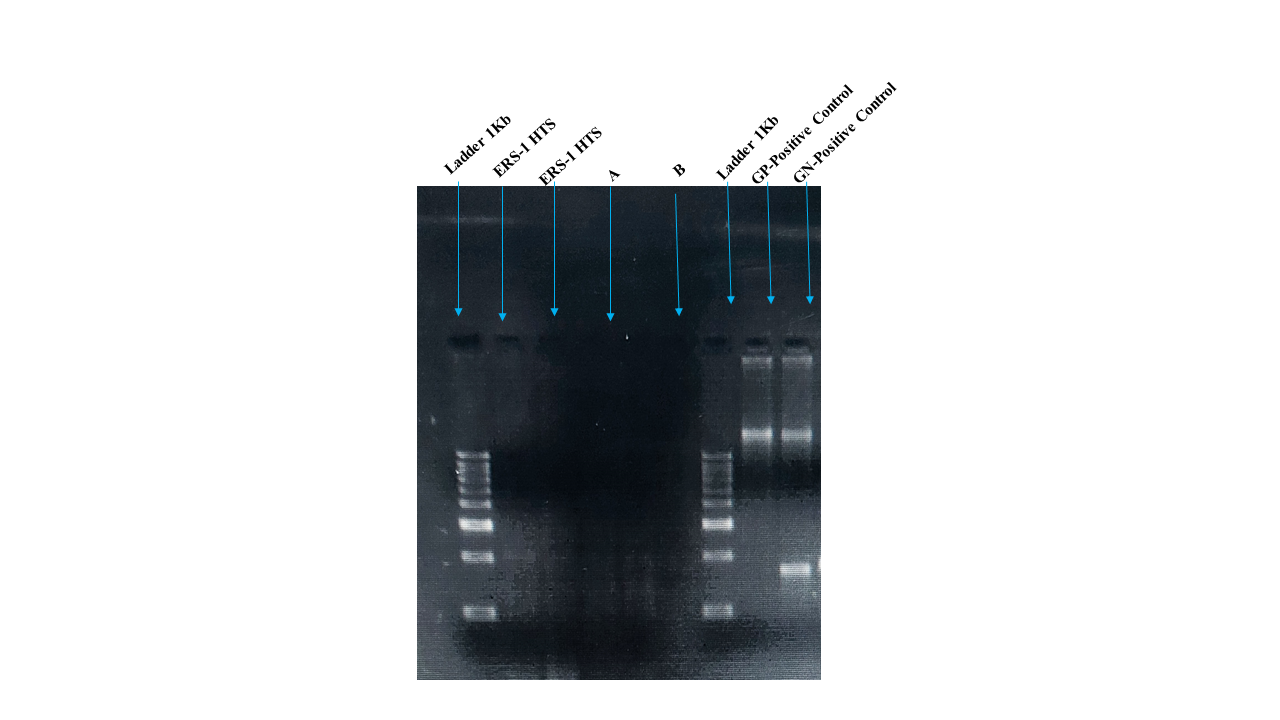

### Fig S2.jpg

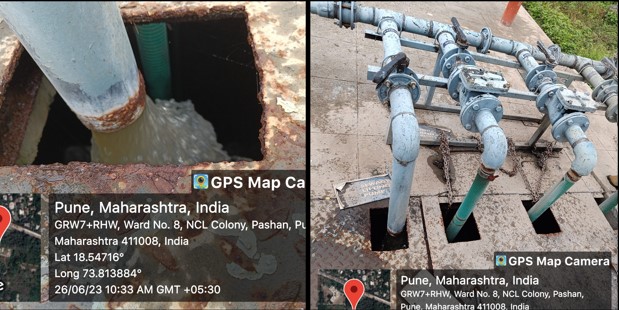

### Fig S2.tif

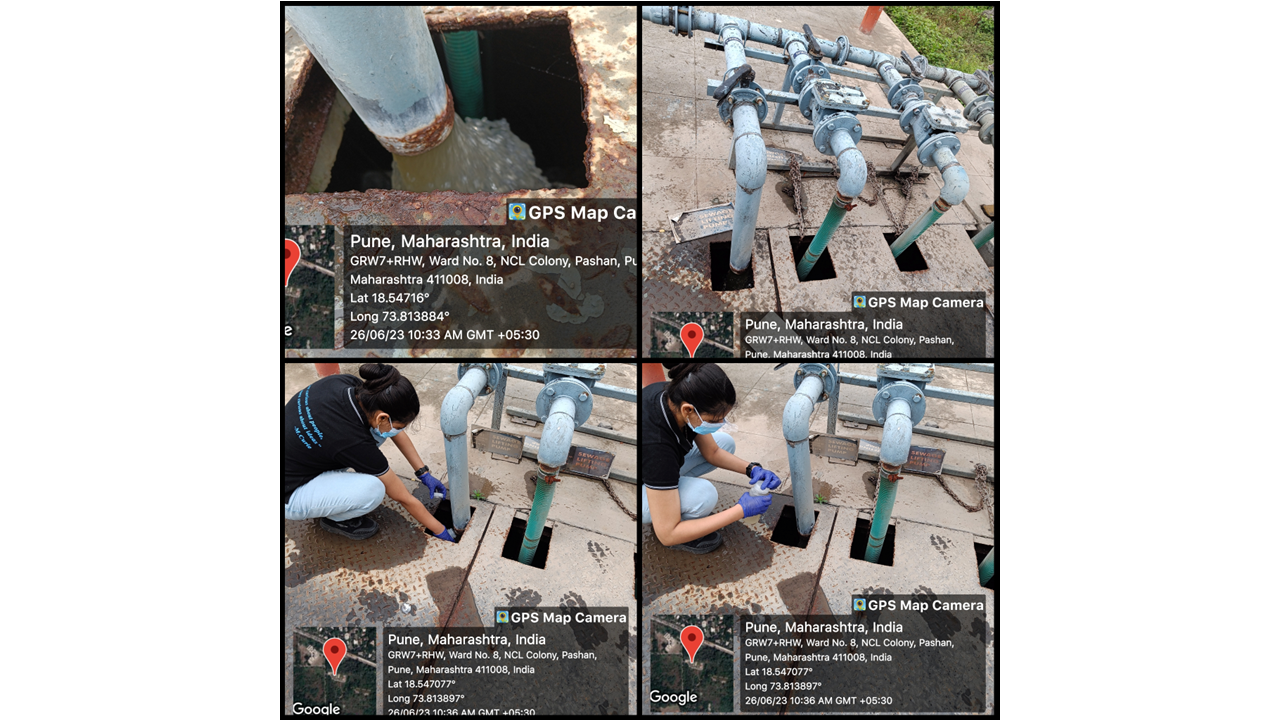
